# Low affinity noradrenergic signaling promotes passive coping during reinforcement behavior

**DOI:** 10.1101/2025.04.05.647394

**Authors:** Ellen M. Rodberg, Elena M. Vazey

## Abstract

During stress and in many psychiatric disorders the ability to appropriately recognize and respond to cues to avoid negative outcomes (e.g., injury) and achieve positive outcomes (e.g., food) is disrupted. A critical component of the stress response is increased noradrenergic tone, acting on lower affinity noradrenergic receptors (α1 and β). Noradrenergic signaling is essential for cue-driven behavior, yet how noradrenergic components of the stress response impacts positive and negative reinforcement behavior is still unknown. We manipulated noradrenergic receptor (α1, α2, and β) activity during an active avoidance and reward seeking task in male and female rats. We found modulating noradrenergic activity on low affinity α1 and β receptors but not α2 receptors disrupted reinforcement behavior. Increased α1, but to a greater extent β receptor activity shifted negative reinforcement behavior away from active avoidance and towards passive coping by increasing decision thresholds. We also found sex differences in the impact of α1 and β noradrenergic activity on reward seeking. Overall, stress related noradrenergic signaling deceased active avoidance behavior by shifting strategy towards passive coping. Reward seeking was more robust in females but not males. Collectively these results identify sex differences in noradrenergic dependent stress response behaviors and highlight low affinity adrenoreceptors as relevant therapeutic targets to mitigate the impact of stress on goal-directed behavior.

**SIGNIFICANCE STATEMENT:** While we know norepinephrine is a fundamental component of the acute stress response, it is unknown how that stress related noradrenergic receptor activity impacts reinforcement behavior, both positive and negative. Here, we show that stress related norepinephrine signaling is differentially involved in positive and negative reinforcement behavior and directly shifts behavioral strategies from active avoidance to passive coping. These results identify that noradrenergic receptor activity can mediate the transition from adaptive to maladaptive behaviors during stress, especially in the context of negative reinforcement.

## INTRODUCTION

Everyday an individual must act based on the relevant cues around them to avoid threats and obtain rewards. These essential goal-directed behaviors are mediated through negative, or positive reinforcement learning. Stress is a major disruptor of goal-directed behavior and reinforcement learning. Stress promotes habitual actions over goal-directed ones and passive coping strategies over active ones. The impact of stress on goal-directed behaviors has been investigated across species using both naturalistic and controlled promotion of physiological stress responses, including pharmacological stressors (Dias-Ferreira et al., 2009; Schwabe and Wolf, 2010, 2011; Schwabe et al., 2011). Many psychiatric disorders show both disrupted goal-directed behavior and heightened stress reactivity. Understanding the mechanisms by which stress can disrupt goal-directed behavior is critical for treating neuropsychiatric diseases from anxiety disorders to substance use disorders (Weinshenker and Schroeder, 2007; Goddard et al., 2010; Naegeli et al., 2018; Keehn et al., 2021).

Stress drives broad and multifaceted physiological responses, including potentiation of noradrenergic signaling with increases in global norepinephrine (NE), and central NE (Valentino and Van Bockstaele, 2008; Uematsu et al., 2017). Central NE signaling is critical for goal-directed behavior, as shown through positive reinforcement learning (Sara and Segal, 1991; Aston-Jones et al., 1997; Bouret and Sara, 2004; Clayton et al., 2004; Aston-Jones and Cohen, 2005a). NE modulates executive functions that underlie goal-directed behavior through a Yerkes Dodson relationship where high NE can drive task disengagement (Aston-Jones and Cohen, 2005b; Arnsten, 2011; Arnsten et al., 2015; Aston-Jones and Waterhouse, 2016; España et al., 2016; Kane et al., 2017; Cope et al., 2019). It is known that heightened noradrenergic tone, such as that elicited by stressors, promotes NE binding on lower affinity, α1 and β receptors both centrally and peripherally (Ramos and Arnsten, 2007). However, whether noradrenergic components of the stress response similarly impact positive and negative reinforcement, and which postsynaptic adrenergic mechanisms are responsible for stress related disruptions in goal-directed behavior remains unclear.

We aimed to understand how noradrenergic components of the stress response, acting on α1 and β receptors, impact goal-directed behavior to avoid threats and/or obtain rewards using female and male rats. We targeted low affinity NE signaling using α1 and β noradrenergic agonists as pharmacological stressors in an active avoidance and reward seeking task. In addition, we used a α2 noradrenergic antagonist to dissociate broad spectrum increases in NE tone from the actions of stress related NE on α1 and β adrenoreceptors. Overall, we found greater resilience to noradrenergic components of the stress response in females. However, across both sexes active avoidance was disrupted by NE acting on β adrenoreceptors. Low affinity NE signaling shifted animals from active to passive coping behaviors on avoidance trials in addition to promoting disengagement from reward seeking in males. Further we found no effect of α2 antagonism on reinforcement behavior which indicates the disruption of goal-directed behavior we saw was not a result of general increased NE tone but specific to low affinity NE targets. Collectively these results identify sex differences in noradrenergic stress response behaviors and highlight low affinity adrenoreceptors as relevant therapeutic targets to mitigate the impact of stress on goal-directed behavior.

## METHODS AND MATERIALS

### Animals

Adult Sprague Dawley rats (n=22; female n=12, males n=10; 6-8 weeks old; 200-300 grams) were purchased from Charles River (Wilmington, MA) and single housed on a reverse 12h light/dark cycle (lights ON at 9:00 PM). Animals had access to ad libitum chow and water until behavioral testing began. During training animals were restricted to ∼80% of ad libitum chow (female=15g, male=20g) until accuracy on the final task was maintained (at least 70% accuracy on all trial types) and then returned to ad libitum access. Weight was monitored throughout experiments to ensure maintenance of at least 80% free fed weight. All animals were handled by an experimenter prior to behavioral training. All procedures were approved by the Institutional Animal Care and Use Committee at the University of Massachusetts Amherst in accordance with the guidelines described in the US National Institutes of Health Guide for the Care and Use of Laboratory Animals (National Research Council 2011).

### Active avoidance and reward seeking task

Operant chambers (Med Associates, St Albans, VT) contained a central well with a recessed panel of LED cue lights and infrared (IR) entry beam, two laterally located retractable levers, a speaker, and grid capable of emitting footshocks. A continuous fan and sound attenuating chambers muffled outside noise. Behavioral data collection was controlled with MedPC IV (Med Associates). All training and tests were performed during the animal’s active cycle in a dimly red lit room.

Animals were trained on an active avoidance and reward seeking trials separately before encountering both trial types in the final task (Figure1A) similar to others previously described (Chowdhury et al., 2019; Capuzzo and Floresco, 2020; Kutlu et al., 2020). First, rats learned to press a lever (right or left, counterbalanced) in response to a specific tone (1kHz) to receive a liquid sucrose reward for reward seeking sessions (Figure 1B, green). Once rats reached criterion (70% accuracy) on reward seeking training, they moved to active avoidance training. During active avoidance training rats learned to press the opposite lever in response to a white noise cue in order to avoid a small footshock (500ms 0.25-0.35mA), (Figure 1B, red). Once animals performed active avoidance trials with 70% accuracy they moved on to the final active avoidance and reward seeking task where avoidance and reward trial types were interleaved within a session. During the final task, rats pressed one lever to receive an appetitive sucrose reward (reward trial), another lever to avoid a series of mild shocks (active avoidance trial), and could press either/no levers press on control trials (neutral trial) in response to distinct auditory cues. Each session consisted of 100 cued trials (40 1kHz reward cue, 40 white noise avoidance cue, and 20 5kHz neutral cue) presented in a pseudorandom order. Each trial began with a variable delay period (mean 20s ± SEM 1.98) after which, a randomly selected auditory cue was presented 5s prior to lever presentation, with the cue persisting until a lever was pressed or the lever presentation period ended (15s), whichever was first. The auditory cues identified which lever should be pressed for positive (receive a reward) or negative reinforcement (avoid foot shocks). Neutral cues require no response, had no consequence and served as a control. Correct lever responses terminated the auditory cue and retracted the response levers in conjunction with a success signal (reward well light) and the appropriate outcome (reward delivery (15% sucrose 95 µl) or shock avoidance/termination). If an incorrect or no lever was pressed within 15s, the trial ended (reward and neutral trials) or an escape period began where the levers remained extended (avoidance trials). Over the 10s escape period, three foot shocks (0.30mA-0.35mA, 500ms) were administered. If the correct avoidance lever was pressed during the escape period, the levers retracted, foot shocks ceased, and the trial ended.

**Figure 1:**
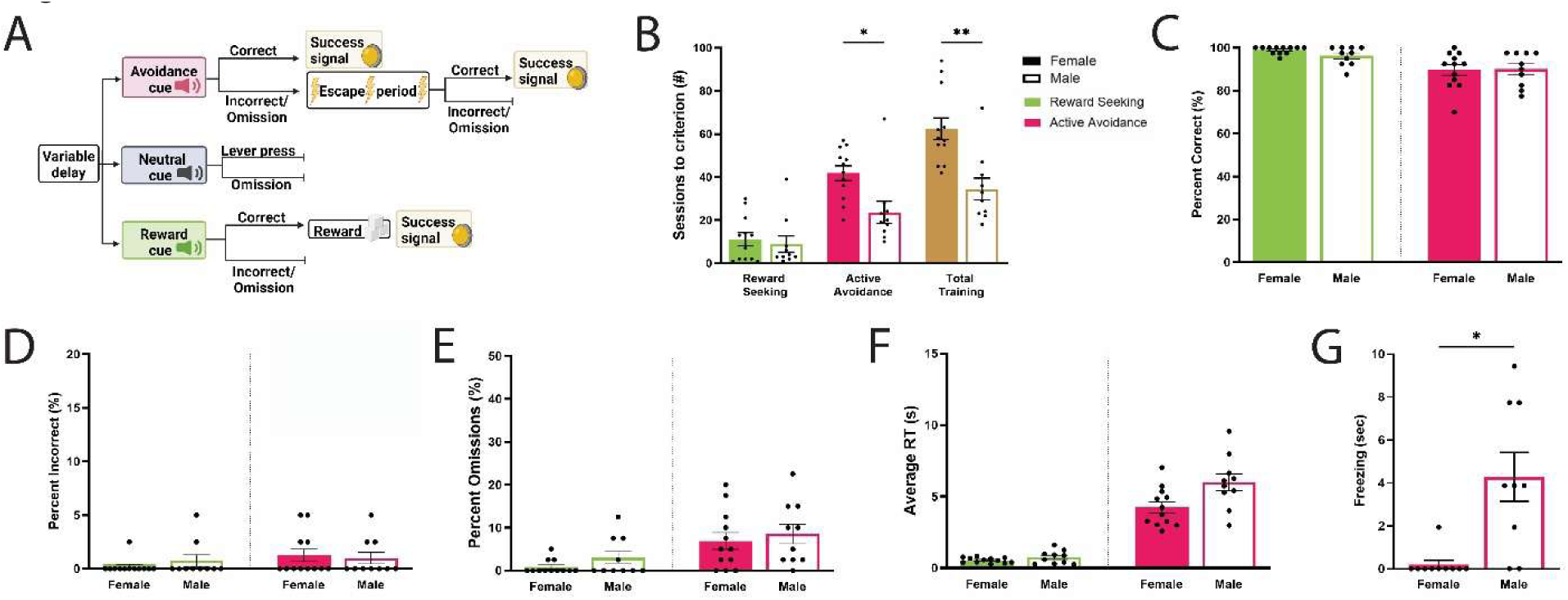
Active avoidance and reward seeking baseline task performance. **A)** Task structure for the active avoidance and reward seeking task with potential outcomes for each trial type. The active avoidance and reward seeking task is comprised of 100 trials; 40 active avoidance, 40 reward seeking, and 20 neutral trials presented pseudorandomly. Each trial began after a variable delay (20s ± SEM 1.98) when rats were presented with one of three auditory cues (<u>avoidance</u>: white noise, top; <u>reward</u>: 1 kHz, bottom; <u>neutral</u>: 5 kHz, middle). Avoidance and reward cues were associated with a correct lever side (right vs. left, counterbalanced across animals) rats learned prior to testing. After 5 seconds of continuous auditory tone, both levers extended while the tone remained on for 15s or until a lever was pressed. If animals pressed the correct lever in 15s they avoid a shock (avoidance trial) or receive a sucrose reward (reward trial) and in both trial types a success signal in the main well would illuminate. If the incorrect lever or no lever was pressed within 15s on a reward seeking trial, no reward was given, and the levers retracted. If the incorrect or no lever is pressed within 15s on active avoidance trials, the escape period was entered. During the escape period, three 500ms (0.30mA-0.35mA) footshocks were administered via the metal grid floor over 10 seconds (3 second break between each shock). If during the escape period the animal pressed the correct lever, the shocks ceased, the levers retracted, and the success signal illuminated. Any response on the neutral trial was not required and received no outcome. **B)** Females took longer to learn the active avoidance and reward seeking task. Training for the final task involved learning the positive reinforcement association (reward seeking) and the negative reinforcement association (active avoidance) separately. Females took longer to reach criterion on the final task (tan, p=0.0036) due to increased sessions to criterion on the negative reinforcement training stage (pink, p=0.0364). Males and females did not differ in their ability to learn positive reinforcement (green). **C-G)** Females and males consistently demonstrate reward seeking and active avoidance behavior after vehicle injections. Across almost all measures of behavioral output, there was no sex differences in baseline vehicle performance during the reward seeking (green) or active avoidance (pink) components of the final task. Females (filled bars) and males (open bars) performed similarly after vehicle administration when comparing **(C)** trial accuracy, **(D)** incorrect responses, **(E)** omitted responses, and **(F)** average reaction times. **(G)** Males displayed significantly more freezing behavior than females during the active avoidance trials (p=0.0258; Kolmogorov-Smirnov test).

### Drugs

After rats could reliably perform the task (>70% accuracy for both positive and negative reinforcement), they were tested with adrenergic compounds at low and high doses in random order, specifically the α1 agonist cirazoline (0.1 mg/kg and 0.3 mg/kg, Tocris, Bristol, UK), β adrenergic agonist clenbuterol (0.005 mg/kg and 0.01 mg/kg; Tocris, Bristol, UK), α2 antagonist atipamezole (0.3 mg/kg and 1 mg/kg; Tocris, Bristol, UK) and 0.9% saline vehicle. Appropriate low and high drug doses were identified from previously published research, cirazoline (Wellman and Davies, 1992; Alsene et al., 2006; Grant et al., 2018), clenbuterol O’Donnell, 1997), and atipamezole (Wright et al., 2012; Grant et al., 2018). Drugs and vehicle were administered IP 30m before testing, at a volume of 1ml/kg. IP injections were chosen as a route of administration to mimic the effects of increased NE signaling that occurs during acute stress –on both central and peripheral receptors. Rats were given at least one day in between injections (vehicle and drug). Drug order was randomized, and each adrenergic agonist day was followed by 1-3 days of baseline performance.

### Analysis

#### Drift Diffusion Model (DDM)

In order to characterize the latent variables underlying decision making we fit single trial reaction time data to a drift diffusion model (DDM) using PyDDM (Shinn et al., 2020). We used PyDDM to fit reaction times on incorrect vs correct trials to a DDM to find values for bound, noise, drift, and non-decision times. We found these variables for each rat for avoidance and reward trials and for each drug treatment separately. The constraints for each model fit were a drift between −10 to 20, noise between .0000001 to 5, bound from .0001 to 20, and nondecision time of 0 to 1, and a poisson mixture with a coefficient of 5, rate of .001. Lastly, we used a dx of .001, dt of .001, and total duration (or maximum reaction time) of 10. The model was then fit, and best parameters reported.

#### DeepLabCut (DLC)

For body part tracking we used DeepLabCut (Mathis et al., 2018) on videos of rats performing the task. We defined the points nose, right eye, left eye, right ear, left ear, head, 5 points down the spine (M1-M5), and tail. The training set was based on 500 labelled frames taken from 20 videos (∼1 per animal). We used aResNet_50 based neural network with default parameters and trained from 200,000 iterations. Randomly assigned 95% of the data used for training and the rest for testing. We then used the X and Y coordinates of key datapoints for behavioral analysis (freezing, movement, etc.), only including those with a cutoff of .75 likelihood for future analysis. **Movement:** Total time (as a percentage) spent moving during the task was found using DeepLabCut coordinates and the DLCanalyzer (Sturman et al., 2020) R script. Movement was calculated using the X and Y coordinates of the body part labelled M1 which was the rats neck, and a movement cutoff of 5 frames and integration period of 5 frames. **Freezing:** We used data generated from DeepLabCut to identify periods of time where the rats were exhibiting freezing behavior, a well characterized fear response. We restricted analysis to time periods during active avoidance trials, from tone onset to levers retracting. Freezing was defined as a lack of head, nose, and body (3 points on spine) movement (less than 1 pixel in X and Y coordinates) for >=1.98s.

#### Leverpress Index

Leverpress index was calculated for neutral trials as shown below for each animal in each condition. The resulting leverpress index of 1 indicates all neutral trial lever presses were on the reward lever and a leverpress index of −1 indicates all neutral trial lever presses were on the avoidance lever.

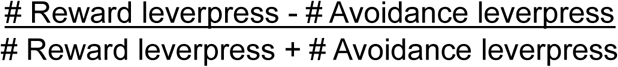

#### Piloerection score

Piloerection score was obtained for each animal after the administration of saline, low doses of cirazoline and high doses of cirazoline. A still image was taken from 30, 60, and 90 minutes into the video recording of the animal performing the task. Images were randomized and scored by an experimenter blind to the condition and animal. Scores given were 0 for no presence of piloerection and 1 for the presence of piloerection. Piloerection score for each animal was the average rating across 30, 60, and 90 minutes.

#### Statistical analysis

Data was analyzed using GraphPad Prism v10 (San Diego, CA), R v4.2.1 (R Core Team (2022), and the Python programming language v3.11.4 (Python Software Foundation). Behavioral data is presented as mean ± SEM after confirming normality (Shapiro-Wilk) unless noted otherwise in the figure legend. Comparisons between groups were made with 2way or REML ANOVA and Sidaks post hoc tests. Graphs were compiled in Prism and figures were compiled in Adobe Illustrator CS (San Jose, CA). Task schematic figure (Figure 1A) was created using BioRender. Bar graphs show the mean ± SEM with data points showing individual animals’ performance. * Indicates a significant effect of sex * (p<0.05); ** (p<0.01); *** (p<0.001); **** (p<0.0001).

## RESULTS

### Female and male rats reliably demonstrate goal-directed behavior for interleaved positive and negative reinforcement

Rats (n=22; female n=12, males n=10) were trained on an active avoidance and reward seeking task (Figure 1A). Animals learned to discriminate stimuli that predicts either positive (leverpress delivers sucrose) or negative (leverpress prevents footshock) reinforcement opportunities. Rats reliably pressed one of two spatially distinct levers in response to the distinct auditory cues (1kHz vs white noise) to receive a sucrose reward (95 µl/trial; reward seeking trials) or avoid a mild footshock (0.3-0.35mA,3 500ms shocks over 10s; active avoidance trials) respectively. Animals also experienced neutral trials (5kHz auditory cue) which were not associated with any reinforcement. Each session contained 40 reward seeking, 40 active avoidance, and 20 neutral trials presented in pseudorandom order.

Rats learned to perform the task by first learning the positive reinforcement association (sessions to criterion: female = 11.17 ± 3.05, male = 8.90 ± 3.77) then the negative reinforcement association (sessions to criterion: female = 41.83 ± 3.45, male = 23.60 ± 5.12). When animals could consistently perform the task (>75% correct on both reward seeking and active avoidance, 3 consecutive days) they moved onto the testing phase. There was a main effect of sex (F _(1, 20)_ = 9.728; p=0.005; 2way ANOVA; Figure 1B), training stage (F _(2.510, 50.19)_ = 80.84; p<0.0001; 2way ANOVA), and interaction (F _(3, 60)_ = 8.267; P=0.0001; 2way ANOVA) on the number of sessions required to learn the task. Females took significantly more sessions to reach testing criterion (p=0.0036), largely due to increased sessions of active avoidance training (p=0.0364). This sex difference aligns with previous studies using a similar positive and negative reinforcement task (Kutlu et al., 2020).

On test days rats received an IP injection (1ml/kg) of either saline vehicle, or an adrenergic based pharmacological stressor. Pharmacological dissociation of adrenergic components of the stress response included a low/high dose of α1 agonist cirazoline (0.01 or 0.03 mg/kg), a low/high dose of β agonist clenbuterol (0.005 or 0.01 mg/kg), or a low/high dose of α2 antagonist atipamezole (0.03 or 1 mg/kg) 20-30 min prior to starting the task. All animals received all doses for within subject comparisons. Drug order was randomized and animals performed several days of baseline performance between test days.

### Baseline reinforcement behavior does not differ across sex

As there were baseline sex differences in the number of sessions required to reach criterion for the final task, we examined if there were any sex differences in reward seeking or active avoidance behavior after vehicle administration. Reward seeking in the task was highly robust during vehicle sessions (female: 98.96% ± 0.48%; male: 96.25% ± 1.36% correct) and rats demonstrated the ability to reliably perform active avoidance behavior after vehicle (females 89.58% ± 2.46%; males 90% ± 2.47% correct). There was no significant difference in reward seeking or active avoidance accuracy between females and males (p=0.6447, p=0.9997, Kolmogorov-Smirnov test; Figure 1C). Once trained, very rarely did a rat press an incorrect lever (Figure 1D). The percent of trials where an incorrect response was made was not significantly different in females or males in either reward seeking (females: 0.21% ± 0.21%, males: 0.75% ± 0.53%; p>0.999, Kolmogorov-Smirnov test) or active avoidance (females: 1.25% ± 0.58%, males: 1% ± 0.55%; p>0.999, Kolmogorov-Smirnov test) trials. Trials where no lever press response was made (omissions) occurred more than incorrect trials but were still uncommon (Figure 1E). Similarly, there was no sex difference in the percent of omitted trials during reward seeking trials (females: 0.83% ± 0.47%; males: 3% ± 1.43%; p= 0.7102, Kolmogorov-Smirnov test) or active avoidance trials (females: 6.88% ± 1.97%; males 8.5% ± 2.30%; p= 0.9981, Kolmogorov-Smirnov test). When a response was made, the average amount of time to press a lever did not differ between female and male rats (Figure 1F) during reward seeking (females: 0.544s ± 0.052s; males: 0.744s ± 0.146s; p= 0.131; Kolmogorov-Smirnov test) or active avoidance trials (females: 4.247s ± 0.389s; males: 5.992s ± 0.59s; p= 0.0738; Kolmogorov-Smirnov test). The only behavioral output that showed a significant sex difference after vehicle administration was the percent of time an animal spent freezing during active avoidance trials (Figure 1G). As expected, females spent significantly less time than males exhibiting freezing behavior between avoidance cue onset and lever press or shock (females 0.193s ± 0.193s; males: 4.27s ± 1.137s; p=0.0258; Kolmogorov-Smirnov test). This is in line with previous research which has identified differences in expression of fear where female rats are more likely to exhibit bolting or darting behavior instead of freezing (Gruene et al., 2015).

### α1 and β adrenergic signaling disrupts positive reinforcement behavior in males but not females

We next examined how the noradrenergic components of stress responses (α1 and β agonism) modulated positive reinforcement, or reward seeking component of the task. When challenged with noradrenergic agonists, reward seeking accuracy decreased (Figure 2A&B) selectively in males with an interaction between sex and drug treatment (F_(4, 80)_ = 7.124; p<0.0001; 2way ANOVA). In males, both low (p=0.0468) and high (p<0.0001) doses of the α1 agonist cirazoline, and high doses of β agonist clenbuterol (p= 0.0031) significantly decreased reward seeking behavior. Compared to vehicle, male positive reinforcement performance after cirazoline decreased by 16.75 ± 8.11% (0.1 mg/kg) and 51 ± 10.79% (0.3 mg/kg). β agonism also showed a modest reduction in reward seeking at the higher dose (0.1 mg/kg) with a 22.75 ± 10.55% reduction in goal-directed behavior. This decreased performance in males was driven not by incorrect lever presses (Figure 2C&D) but rather primarily mediated by an increase in omitted responses (Figure 2E&F) where no leverpress was made (cirazoline low p=0.0433, cirazoline high p<0.0001, clenbuterol high p=0.0036). In contrast to the decrease in positive reinforcement behavior in males, accuracy on reward seeking trials in females was relatively stable and they maintained a high level of reward seeking under all doses of α1 and β agonists (p>0.05). We did, however, see a shift in decision dynamics in females, indicating strategy adjustments in the face of stress related adrenergic potentiation.

**Figure 2:**
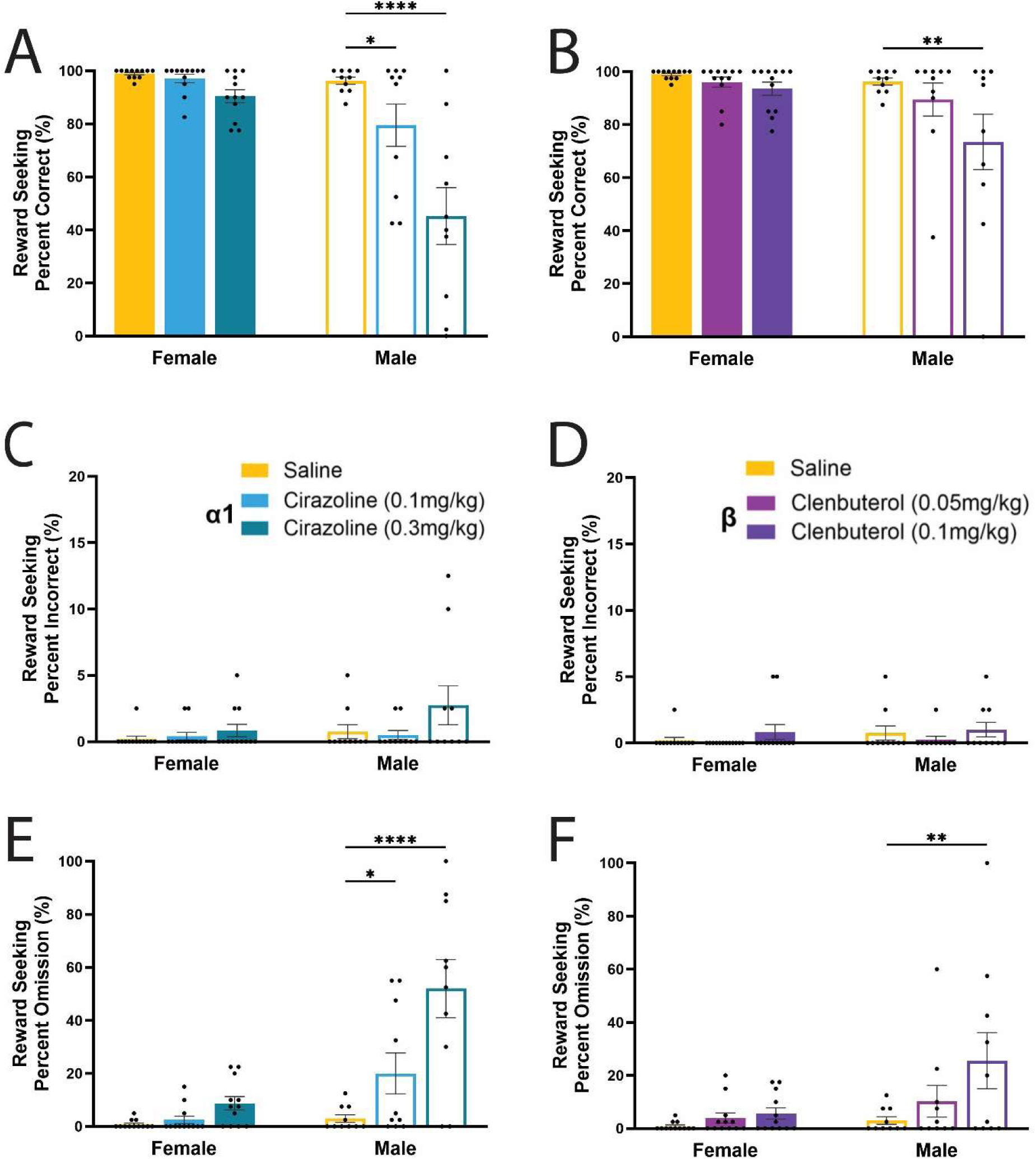
Reward seeking in males was sensitive to stress related increases in NE tone. **A-B)** Accuracy on reward seeking trials decreased in males (unfilled bars) but not females (filled bars) after α1 (**A**, blue, left) and β (**B**, purple, right) agonism. The percent of trials where a correct response was made decreased in males after both doses of cirazoline (low p=0.0468, high p<0.0001) and high doses of clenbuterol (p= 0.0031). **C-D)** Incorrect lever press responses on reward seeking trials remained low and did not increase after α1 **(C)** or β **(D)** agonism. **E-F)** In males, cirazoline (**E**, low p=0. 0433, high p<0.0001) and high doses of clenbuterol (**F**, p=0.0036) increased the percent of omitted trials where no lever press response was made. Females did not increase omitted reward seeking trials after α1 or β agonism.

### Females persist in reward seeking but slow information accumulation with heightened NE tone

Reaction time, measured as the amount of time for an animal to respond with a lever press once the levers extend for each trial, was increased by noradrenergic pharmacological stress (Figure 3A&B; F_(4,77)_=10.80; p<0.0001; REML ANOVA) with a sex and treatment interaction (F_(4,77)_=4.146; p=0.0043; REML ANOVA). Reaction times significantly increased in females after high doses of cirazoline (p=0.0010, 1.458s ± 0.234s) and low doses of clenbuterol (p=0.0383, 1.135s ± 0.211s) compared to vehicle (0.543s ± 0.052s). Cirazoline (α1 agonist) but not clenbuterol (β agonist), significantly slowed male responses on positive reinforcement trials, increasing reaction times after both doses (low p=0.0126, 1.488s ± 0.251s; high p<0.0001, 1.943s ± 0.305s) compared to vehicle (744.1 ± 145.8ms).

**Figure 3:**
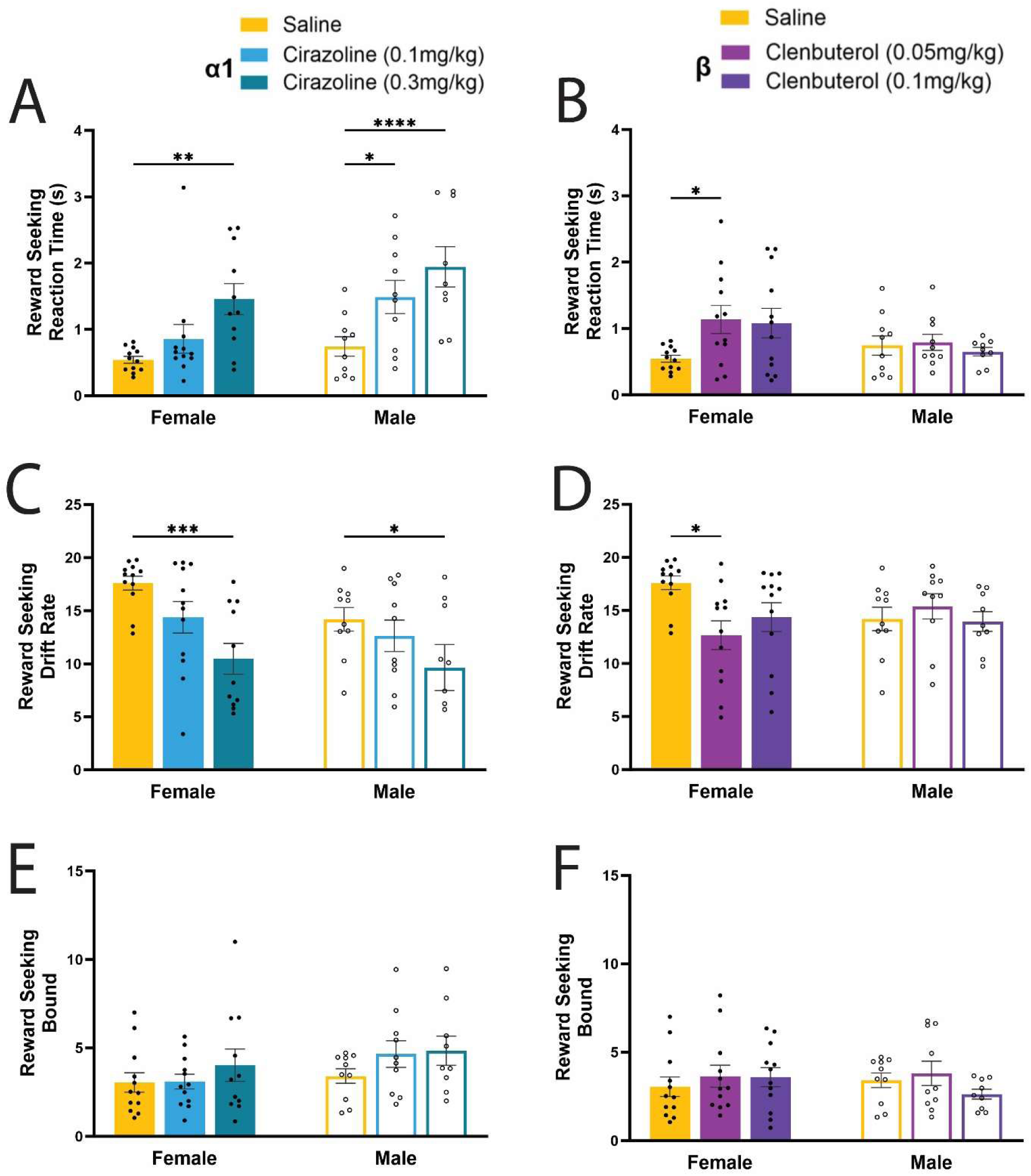
The noradrenergic component of stress decreased the rate of evidence accumulation on reward seeking trials, resulting in increased reaction times. **A-B**) The amount of time for a lever press response to be made on reward seeking trials increased after high doses of (**A**) cirazoline (p=0.0010) and low doses of (**B**) clenbuterol (p=0.0383) in females. In males, both doses of cirazoline (low p=0.0126, high p<0.0001) increased reaction times. **C-D**) Using the drift diffusion model (DDM) to characterize the latent variables underlying decision making, drift rate, or the rate of evidence accumulation for a decision to be made, decreased after high dose of (**C**) cirazoline (females p= 0.0002, males p=0.0455). Drift rate also decreased in females after low doses of (**D**) clenbuterol (p=0.0110). **E-F**) The boundary separation parameter of the DDM, which represents the amount of time/evidence needed for a decision to be made, was not different in males of females after (**E**) α1 or (**F**) β agonism.

To further characterize the latent processes underlying decision-making and the shift in reaction times during positive reinforcement behavior, we implemented a drift diffusion model.

We found a main effect of treatment in the drift rate (Figure 3C&D), or the rate of information accumulation (F_(4,77)_=5.925; p=0.0003; REML ANOVA). In females, how quickly evidence accumulated in favor of a particular response was significantly decreased after high doses of cirazoline (p= 0.0002) and low doses of clenbuterol (p=0.0110). In males, drift rate was significantly decreased only after high doses of cirazoline (p= 0.0455). Other DDM variables, such as bound/boundary separation parameter, which reflects the amount of time/evidence needed for decision making (speed-accuracy tradeoff) were not altered after noradrenergic modulation (Figure 3E&F).

Together, this data suggests that increased low affinity NE signaling, similar to that elicited by stress, drives different adaptations in reward seeking behavior from females and males. Specifically, noradrenergic stressors slow information accumulation for positive reinforcement but females continue to successfully seek rewards, whereas males disengage from reward seeking. Although NE α1 receptor stimulation slowed decision times in males they primarily disengaged from positive reinforcement behavior as shown by increased omissions (Figure 2F). β adrenergic agonists also promoted disengagement in males but did not shift decision dynamics. This disengagement from males is in stark contrast to females that persisted in goal directed behavior for positive reinforcement despite slowed accumulation of evidence necessary for successful choices.

### Stress related increases in noradrenergic tone severely impacted active avoidance

We next examined the impact of noradrenergic agonists on negative reinforcement trials where rats had to press a lever to avoid a mild foot shock, obtained from the same sessions in which we measured reward seeking. Unlike positive reinforcement (above), goal directed negative reinforcement behavior in both sexes was significantly impacted by noradrenergic stressors. Noradrenergic agonists (α1 & β) significantly reduced avoidance behavior (Figure 4A&B; F_(4, 80)_ = 20.86; p<0.0001; 2way ANOVA) with a treatment and sex interaction (F_(4, 80)_ = 2.830; p=0.0299; 2way ANOVA) where males and females were negatively impacted by all drugs and doses except for female’s resilience to low doses of cirazoline. After low and high doses of the α1 agonist cirazoline, which also disrupted reward seeking in males, negative reinforcement behavior decreased by 26.75% ± 9.32% (p=0.0048) and 62% ± 9.70% (p<0.0001), respectively. Unlike positive reinforcement behavior, females showed a significant 35.42% ± 9.21% (p<0.0001) decrease in accuracy on negative reinforcement trials after high dose cirazoline. The response to β NE agonist clenbuterol, at both doses, was similar for both sexes showing reduced accuracy on active avoidance trials. Males showed a reduction of 33% ± 7.19%, (p=0.0014) and 45.75% ± 10.65%, (p<0.0001), after low and high dose clenbuterol respectively. In females, performance after low and high dose clenbuterol dropped by 46.04% ± 12.41% and 40.63% ± 9.84% (both p<0.0001).

**Figure 4:**
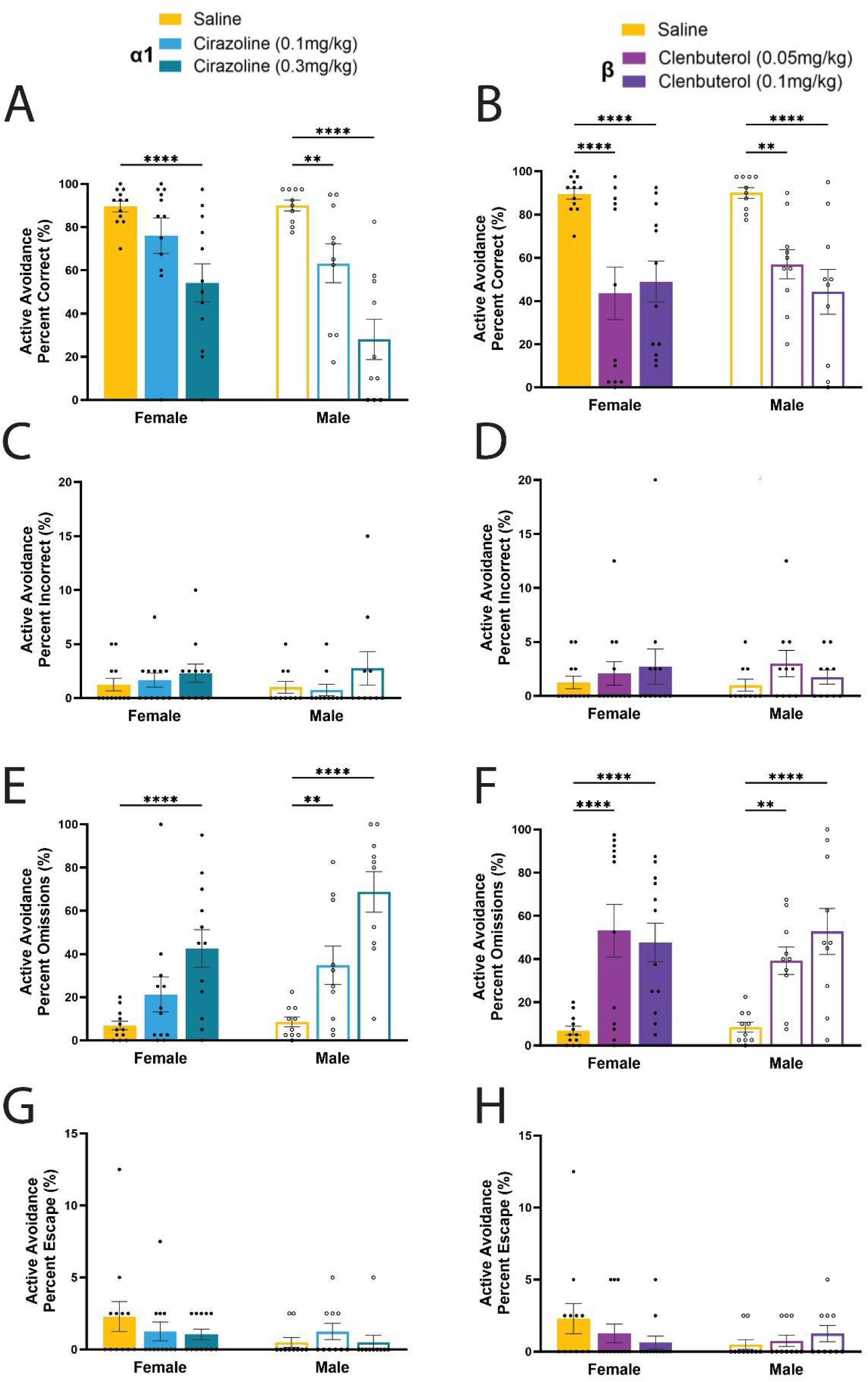
Active avoidance behavior was highly sensitive to stress related noradrenergic modulation. **A&B**) Accuracy on active avoidance trials decreased after all doses of α1 agonist cirazoline (**A**) (female: high p=0.0003; male: low p=0.0184, high p<0.0001) and β agonist clenbuterol (**B**) (female: low p<0.0001, high p<0.0001; male: low p= 0.0022, high p<0.0001) except for low doses of cirazoline in females. **C&D**) Incorrect responses did not change after (**C**) α1 or (**D**) β agonist administration in males or females. **E&F**) Decreased accuracy on active avoidance trials was a direct result of increased omissions, trials where no lever press response was made, after (**E**) α1 (female: high p =0.0002; male: low p=0.0213, high p<0.0001) and (**F**) β agonism (female: low p<0.0001, high p<0.0001; male: low p= 0.0049, high p=<0.0001). **G&H**) Escapes, trials where a correct response is made during the shock period which would extinguish the remaining shocks, did not change after (**G**) α1 or (**H**) β agonist administration in males or females.

As with reward seeking, this shift in performance was not a result of incorrect responses (Figure 4C&D) but a result of increased omitted responses (Figure 4E&F). There was a main effect of treatment (F_(4, 80)_ = 20.04; p<0.0001; 2way ANOVA) and a sex and treatment interaction (F_(4, 80)_ = 2.854; p=0.0289; 2way ANOVA) on omissions during active avoidance trials. Compared to vehicle, females increased omissions after high doses of α1 cirazoline (p=0.0002) and both doses of β agonist clenbuterol (p<0.0001 for each). In males, all doses of noradrenergic agonists increased omissions (cirazoline: low p=0.0213, high p<0.0001; clenbuterol: low p= 0.0049, high p=<0.0001).

Avoidance trials provided an opportunity for rats to demonstrate reactive escape behavior by lever pressing during the “escape period” - the 10s window in which shocks were administered. This behavior was only seen on a small number of trials. We did not find any difference in escape behavior based on sex or treatment (Figure 4G&H). If the rats did not decide to initiate a response prior to the foot shocks, noradrenergic agonism did not alter the probability of responding during the escape period.

This behavioral profile suggests that after administration of noradrenergic agonists, the pharmacological stressor shifts behavioral strategies towards a learned helplessness like phenotype where behavioral responses to avoid negative outcomes are omitted and instead animals display passive coping.

### Noradrenergic agonists increase reaction time and decision thresholds for negative reinforcement

Although negative reinforcement behavior was reduced with enhanced noradrenergic signaling, approximately half the trials were still completed. We investigated the response characteristics of behavior on active avoidance trials where a response was made. We found reaction times increased as a main effect of treatment (Figure 5A&B) (F_(4, 75)_ = 9.663; p<0.0001; REML ANOVA), with interactions between treatment and sex where male reaction times were significantly increased after high doses of both cirazoline and clenbuterol and female reaction times were only impacted by clenbuterol (F_(4, 75)_ = 3.010; p= 0.0233; REML ANOVA). Compared to vehicle (female: 4.201s ± 0.396s, male: 5.974s ± 0.590s) cirazoline increased reaction times in males at high doses (8.267s ± 1.073s, p= 0.0155). Female reaction times where not significantly changed by α1 neuromodulation. However, clenbuterol increased reaction time in both females (low: 8.316s ± 1.33s, p<0.0001; high: 6.677 ± 0.814s, p= 0.0051) and males (high dose only: 8.329s ± 0.605s, p= 0.0118).

**Figure 5:**
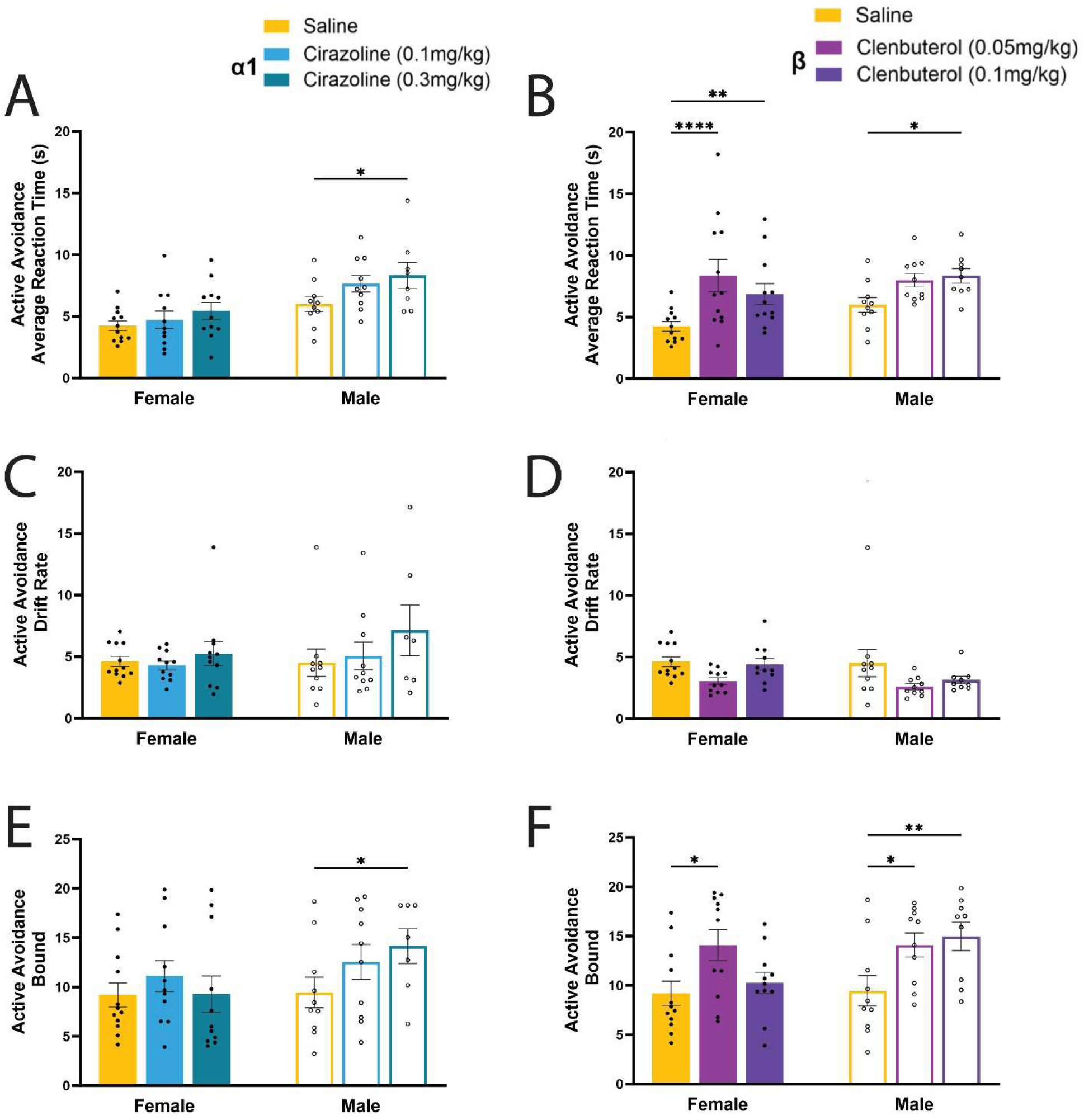
Noradrenergic stress increased reaction times and boundary separation during decision making on active avoidance trials. **A&B**) The average time to respond to an active avoidance trial increased in males after high doses of (**A**) cirazoline (p=0.0155). (**B**) Clenbuterol increased reaction times after high doses in males (p=0.0118) and both doses in females (low p<0.0001, high p=0.0051). **C&D**) Drift rate, a component of the DDM related to evidence accumulation, was not greatly impacted by noradrenergic stressors. α1 and β NE agonism resulted in a main effect of treatment on the drift rate during active avoidance behavior. **E&F**) Bound, a component of the DDM that represents the time/amount of evidence that must be accumulated to respond to a stimulus was impacted by NE modulation. (**E**) High doses of cirazoline increased bound in males (p=0.0204). (**F**) Clenbuterol had a much larger effect increasing bounds in males at both doses (low p=0.0368, high p= 0.0068) and in females at low doses (p=0.0139).

We again used the drift diffusion model to characterize the latent variables underlying decision making changes caused by noradrenergic component of stress (Figure 5C-F). Unlike during reward seeking, drift rate showed limited impact of α1 or β NE modulation during active avoidance, although there was a main effect of treatment (Figure 5C&D; F_(4, 92)_ = 4.643; p= 0.001; REML ANOVA) no post-hoc differences between groups were found. However, we found strong evidence for an increase in the threshold needed to make a choice with a main effect of treatment (Figure 5E&F; F_(4, 72)_ = 4.317; p=0.0035; REML ANOVA) on the decision bound. As with reaction times, we found an increased bound variable in males after high doses of cirazoline (p=0.0204) but no change in females. In females, low doses of clenbuterol increased the bound (p=0.0139) and in males both the low (p=0.0368) and high (p=0.0068) doses of clenbuterol increased bound. These results highlight that heightened NE tone can mediate different cognitive demands for positive and negative reinforcement during decision making across sexes.

### NE agonists shift behavior away from goal directed avoidance and toward passive coping in females

After α1 & β NE agonist administration, negative reinforcement behavior was significantly impaired, and decision thresholds increased. We investigated whether a shift to passive behavioral coping strategies was underlying decreased active avoidance performance. Freezing is a well-defined fear response that can be driven by footshocks and the cues that predict them. We found a main effect of treatment (Figure 6A&B; F_(4, 68)_ = 6.764; p= 0.0001; REML ANOVA) on the amount of time spent freezing during avoidance trials. Females significantly increased the time spent freezing after high doses of the β agonist clenbuterol (episodes >1.9 s; p=0.0006). Reciprocally, when comparing the total percentage of time spent moving during the task, we found that clenbuterol, but not the α1 agonist cirazoline, decreased the percentage of time moving (Figure 6C&D). We found a main effect of treatment (F_(4, 72)_ = 24.15; p<0.0001; 2way ANOVA) and a sex and treatment interaction (F_(4, 72)_ = 3.911; p=0.0063; 2way ANOVA). High doses of clenbuterol in males (p=0.0279) and females (p<0.0001) as well as low doses of clenbuterol in females (p<0.0001) decreased the total percentage of time spent moving over the entire session. While cirazoline had no impact on movement, we did notice an increased prevalence of piloerection after cirazoline administration (F _(1.303, 23.45)_ = 18.83; p<0.0001; 2way ANOVA), a known impact of this drug ((Alsene et al., 2006) and increased α1 tone (Supplemental Figure 1). These results show that β NE receptor activation promotes passive freezing behavior over active avoidance strategies.

**Figure 6:**
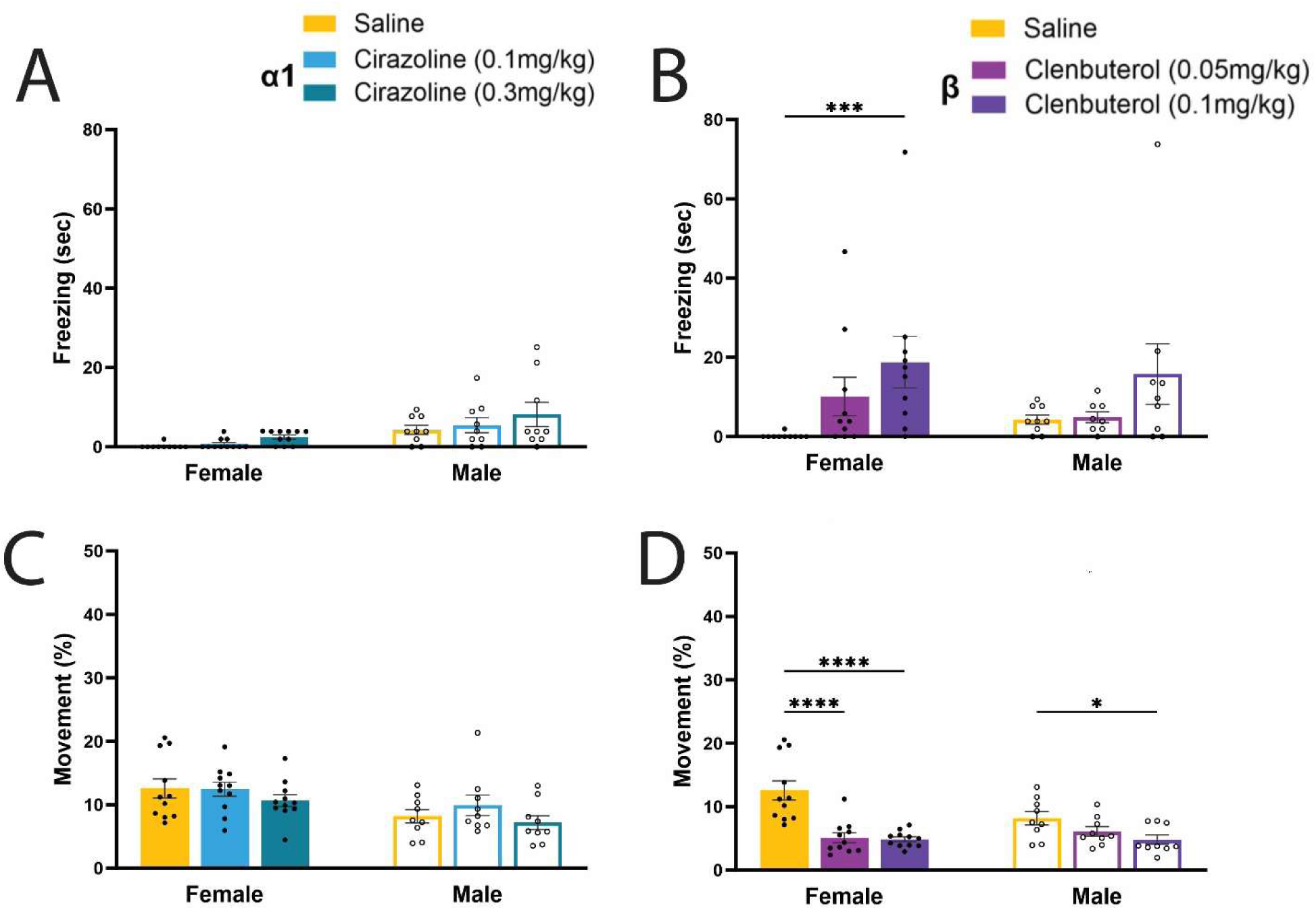
Clenbuterol decreased movement and increased freezing, particularly in females. **A&B**) (**A**) Cirazoline had no impact on the amount of time freezing. (**B**) In females, high doses of clenbuterol increased the amount of time spent freezing during active avoidance trials (p=0.0006). **C&D**) (**C**) The percentage of time spent moving across the entire task was not impacted by cirazoline. (**D**) Clenbuterol, however, decreased the total percentage of time spent moving during the active avoidance and reward seeking task in females (low p<0.0001, high p<0.0001) and males (high p=0.0279).

### Neutral trials reveal a propensity to prioritize safety over reward, especially in females, that is disrupted by increased β NE activity

We looked at behaviors during neutral trials to determine whether subjects showed a prepotent behavioral bias. On neutral trials, rats hear a distinct tone and can press either lever but there is no incentive or punishment for doing so, they receive no reward or shock regardless of their action. During vehicle sessions we found a sex difference in the propensity to respond on a neutral trial where females lever pressed on the majority of neutral trials whereas males did not (Figure 7A; p=0.0025; Mann Whitney test; 67.5% vs. 35%).

**Figure 7:**
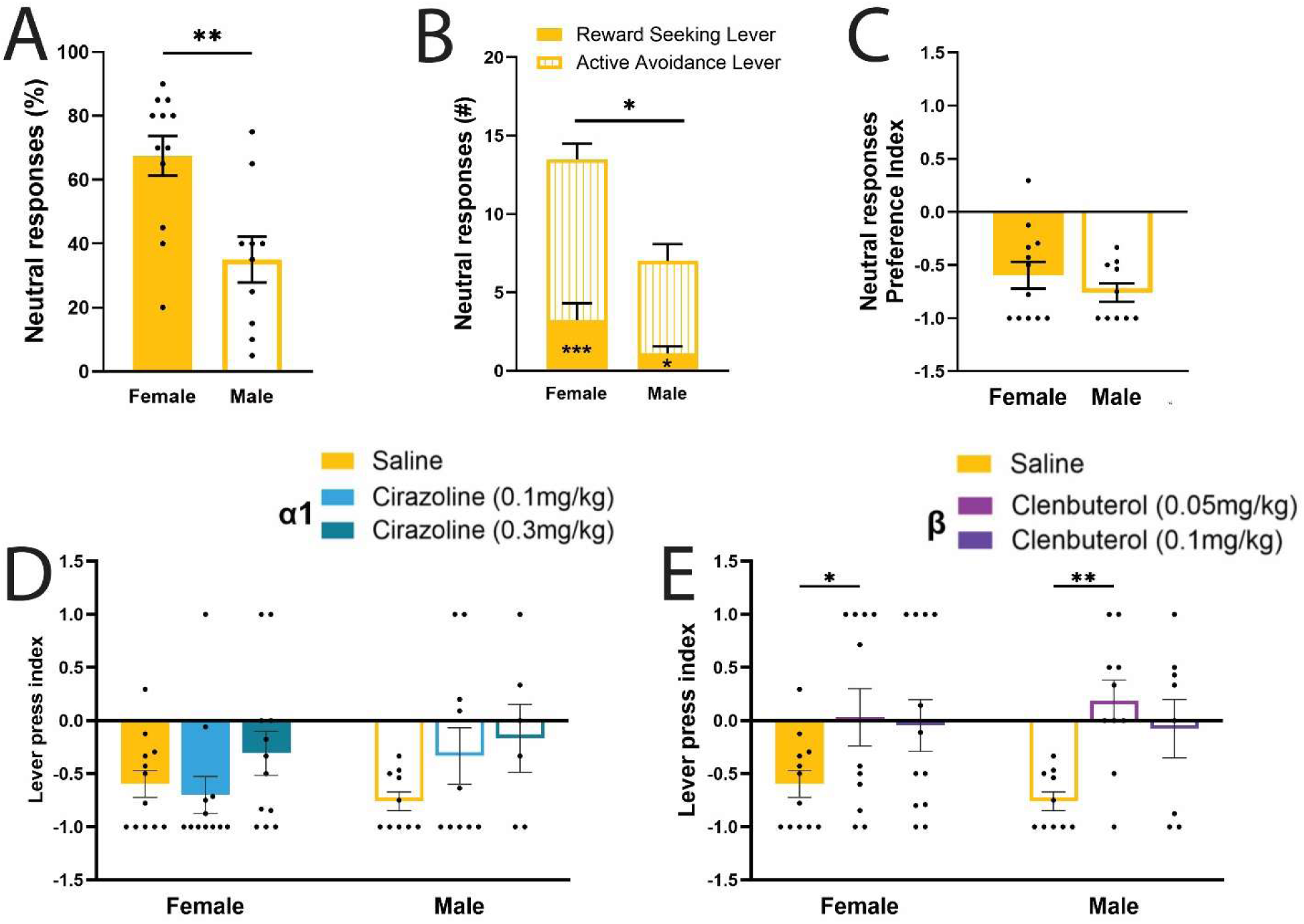
β agonism shifts neutral trial strategy. **A)** Strategy after saline administration shows both sexes lever press on neutral trials but females respond to a higher percentage of neutral trials than males (p=0.0025; Mann Whitney test; 75% vs. 37.5%). **B)** Further dissecting the neutral trial responses after saline revealed that female (p=0.0001) and male (p=0.0133) rats press the active avoidance assigned lever (striped bar) more than the reward seeking assigned lever (solid bar). Furthermore, there was a sex difference where females pressed the avoidance lever more than males (p=0.0106). **C)** Almost all rats after saline display a strong preference for pressing the active avoidance lever during neutral trials. Lever press index takes the number of presses on the reward lever minus the avoidance lever divided by the total neutral trial presses to normalize the total presses across animals. Lever press index closer to −1 indicates more active avoidance lever presses during neutral trials and a lever press index closer to 1 indicates more reward seeking lever presses during neutral trials. **D&E**) Low doses of clenbuterol shift the strategy implemented during neural trials away from active safety. Clenbuterol (**E**) but not cirazoline (**D**) shifted the leverpress index in a more positive direction in males (p=0.0014) and females (p=0.0346).

We investigated which lever animals chose on neutral trials - the avoidance or reward lever. We found an overall higher propensity to press the active avoidance lever compared to the reward seeking lever on neutral trials, that was especially prevalent in females (Figure 7B). We found a main effect of sex (F_(1, 20)_ = 11.85; p=0.0026; 2way RM ANOVA) and a main effect of lever type (F_(1, 20)_ = 36.55; p<0.0001; 2way RM ANOVA). Post hoc tests revealed that females pressed the avoidance lever more than males (p=0.0106). Furthermore, within each sex, rats pressed the avoidance lever more than the reward lever (females p=0.0001, males p=0.0133). This illuminates an overall strategy to prioritize safety and a sex difference where females are more likely to respond on neutral trials and respond more on the avoidance lever.

To account for sex differences in overall probability of neutral responses we normalized the data using a lever press index. The lever press index subtracts the number of presses on the avoidance lever from the reward lever and divides this by the total neutral trial lever presses to obtain one value for each animal. If this index was closer to 1 then the rat was more likely to press the reward lever during neutral trials and if the index was closer to −1 the rat was more likely to press the avoidance lever on neutral trials. Observing the leverpress index after vehicle confirmed evidence for a prepotent strategy in both sexes to prioritize safety by pressing the avoidance lever (Figure 7C).

Interestingly, we found that noradrenergic treatment shifted the lever press index on neutral trials (F_(4, 73)_ = 6.917; p <0.0001; REML ANOVA; Figure 7D&E). Specifically, low doses of clenbuterol shifted the leverpress index in a more positive direction (preferring reward seeking) in males (p=0.0014) and females (p=0.0346). This reveals that although there is no incentive to press any lever during neutral trials, animals are likely to press the avoidance lever in what is presumably a safety strategy. However, β NE agonism disrupts this active safety strategy and promotes more learned helplessness even outside avoidance trials.

### General increased noradrenergic tone did not alter behavior

Lastly, we tested if the impacts on active avoidance and reward seeking identified above were a result of a general increased noradrenergic tone vs. increased NE action on lower affinity α1 & β receptors involved in the stress response. To do this, we tested the same animals with low and high doses of α2 antagonist atipamezole. α2 antagonists acting presynaptically on LC neurons promote NE release by decreasing autoinhibition (Lapiz et al., 2007). We found no dose of atipamezole altered reward seeking or active avoidance trials in females or males (Figure 8). This highlights that the deficits on active avoidance and reward seeking behavior, strategy shift from active to passive coping, were specific to stress related noradrenergic signaling via α1 and β NE receptors.

**Figure 8:**
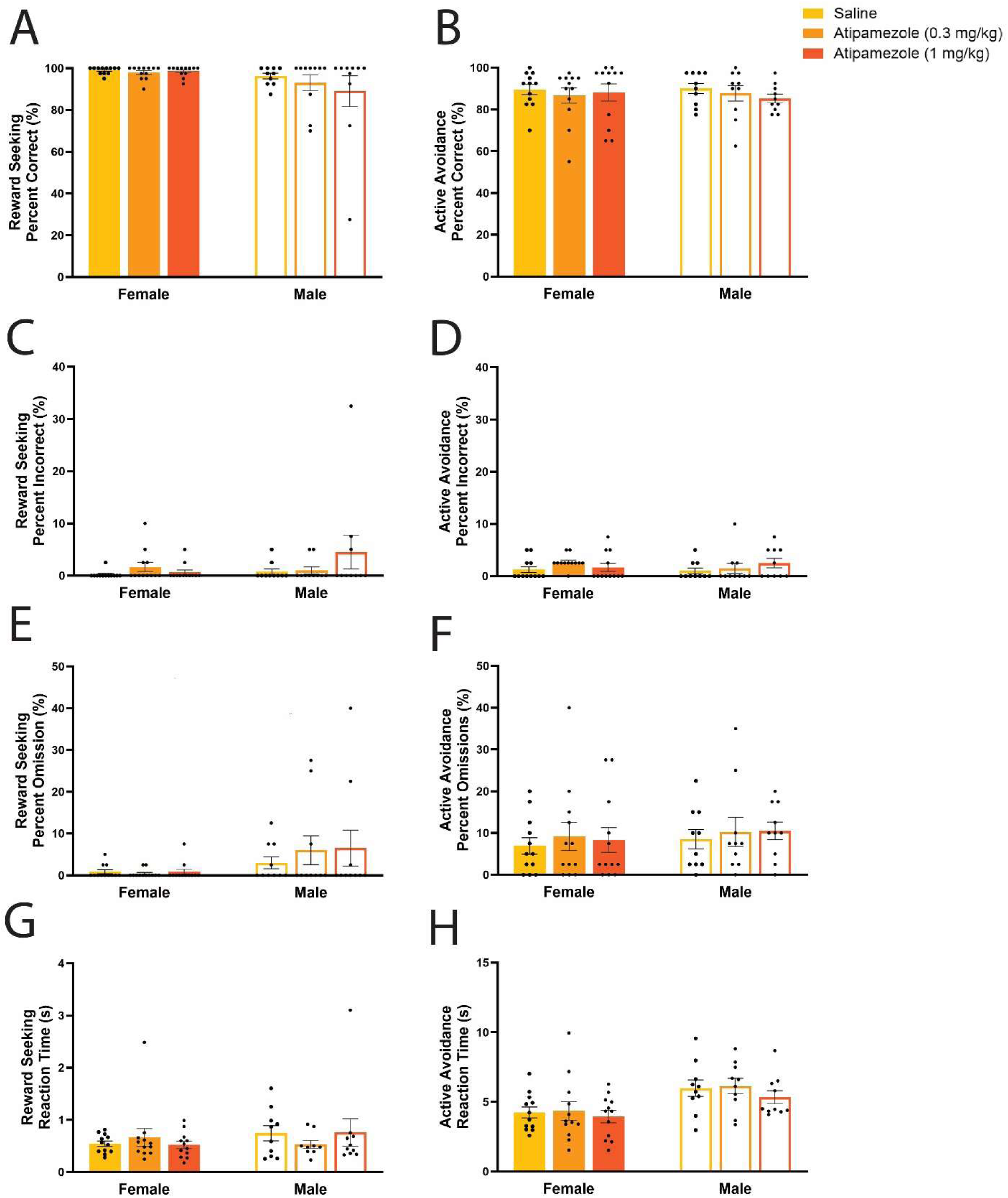
α2 antagonism did not alter active avoidance or reward seeking behavior. Increasing general noradrenergic tone by antagonizing α2 did not impact reinforcement behavior. **A&B)** No dose of atipamezole significantly altered accuracy for (**A**) reward seeking or (**B**) active avoidance. **C&D)** α2 antagonism did not impact the percent of incorrect responses during (**C**) reward seeking or (**D**) active avoidance trials. **E&F)** The percent of omitted trials during (**E**) reward seeking and (**F**) active avoidance was unchanged after atipamezole administration. **G&H)** Reaction times during (**G**) reward seeking and (**H**) active avoidance trials were not impacted by α2 antagonism.

## Discussion

We identified both independent and sex dependent roles for stress related noradrenergic signaling on reinforcement behavior. We found stress-like NE signaling at low affinity adrenoreceptors, particularly β adrenoreceptors, reduced active avoidance behavior and promoted passive coping strategies during negative reinforcement in both males and females. In addition activation of low affinity α1 and β noradrenergic signaling promotes disengagement of reward seeking behavior in males. Despite reduced information accumulation rates, females persisted in positive reinforcement behaviors under stress-like activation of NE receptors, showing a distinct behavioral response from males.

### Noradrenergic activation differentially impacts reward seeking highlighting stress resilience phenotypes

We found that males, but not females, were sensitive to NE modulation during reward seeking, increasing the percent of trials where no response was made. α1 agonism promoted more behavioral disengagement than β agonism during male reward seeking (Figure 2), while general increases in NE tone with α2 antagonism did not (Figure 8). We, and others have previously shown that central LC noradrenergic activation is sufficient to promote task disengagement for positive reinforcement in males (Kane et al., 2017; Cope et al., 2019). When males did participate in reward seeking, α1 agonism increased reaction times and decreased evidence accumulation rates (drift component of DDM). This may suggest α1 and β are differentially involved in positive reinforcement behavior, α1 decreasing evidence accumulation in males, and β shifting the behavioral strategy towards disengagement.

Unlike males, females maintained reward seeking under NE modulation despite increased effort reflected in reaction times and decreased drift rates. These differences may be driven by innate differences in central NE size, organization, and responsivity to stressors or regional sex differences in adrenergic receptor distribution in PFC and other regions that we and others have identified previously (Ohm et al., 1997; Pinos et al., 2001; Bangasser et al., 2011, 2016; Rodberg et al., 2023).

### Low affinity norepinephrine signaling induces passive coping strategies for negative reinforcement

We identified sex differences during training where females took longer to initially engage in active avoidance behaviors, confirming a pattern previously identified in mice (Kutlu et al., 2020). Despite the difference in initial training both sexes robustly maintained active avoidance behaviors until challenged with noradrenergic agonists.

In contrast to our findings during reward seeking, active avoidance behavior was negatively impacted by α1 and β agonists in both sexes. All treatments except low doses of cirazoline in females promoted omissions during negative reinforcement trials. This suggests that NE signaling promotes a switch in behavioral strategy from active avoidance/proactive coping to reactive/passive coping. Instead of proactively preventing the shock, as they were highly trained to do, rats disengaged and treated the negative outcome as unavoidable with reduced avoidance and escape behavior. This shift was reflected in an increased decision threshold in both sexes. We also saw general movement disruptions during active avoidance trials with increased periods of freezing in response to active avoidance cues in females after β agonism.

Stressors and negative reinforcement can elicit active vs. passive coping by engaging different neural circuits. Active coping occurs in response to escapable stressors (such as during active avoidance trials), however active responses must often be learned, as was the case in our study. Innate responses to aversive events are usually passive behaviors e.g. freezing (Maier and Seligman, 2016). Passive behaviors continue when the aversive stimulus is unavoidable resulting in learned helplessness. However here, active avoidance strategies reverted to passive and learned helplessness like behaviors when low affinity noradrenergic receptors were stimulated. Multi-faceted passive behaviors emerged including freezing, increased reaction times, increased omissions and shifting decision dynamics with increased DDM bounds and decreased drift rates. These changes provide evidence that as part of a physiological stress response α1 and β noradrenergic activation drives the emergence of passive coping strategies in response to negative stimuli and identify heightened noradrenergic responses as a mechanism that may predispose individuals to learned helplessness.

Active coping requires engagement of prefrontal cortex (PFC) outputs to regulate several targets. Even learned helplessness can be overcome by engaging appropriate medial PFC outputs, such as those to the dorsal raphe (Baratta et al., 2009; Warden et al., 2012). Within the PFC, norepinephrine is tightly involved in neural representations of stimuli (Foote et al., 1975; Berridge and Waterhouse, 2003; Ghosh and Maunsell, 2024) as well as the maintenance and execution of action plans (Bornert and Bouret, 2021). NE does this in part by regulating working memory (α1 and α2) and longer-term conditioned fear memories (β) (Murchison et al., 2004; Ramos and Arnsten, 2007; Berridge and Spencer, 2016; Giustino and Maren, 2018). Altering NE signaling in the PFC can disrupt neural representation of relevant stimuli (Ramos and Arnsten, 2007; Rodberg et al., 2023). Stress is known to disengage PFC processing of relevant information (Arnsten, 2015; Datta and Arnsten, 2019). Changes in neural representation and memory processing in the PFC or relevant PFC outputs for those signals via increased α1 and β NE tone in the current study is a likely driver of passive responses to negative reinforcement stimuli.

Outside the PFC, active coping engages paragigantocellularis enkephalin neurons (PGi-enkephelin) inputs to LC to promote resilience to stress related effects. In contrast passive coping engages input from CRF positive central amygdala neurons (CNA-CRF) neurons to LC, resulting in feedforward LC hyperactivity and enhanced sympathetic drive (Reyes et al., 2015; Wood and Valentino, 2017) We identified that the active/proactive coping behaviors engaged at baseline switched to passive/reactive after administration of α1 and β agonists, the NE pharmacological stressors could mimic LC response elicited by a shift in regulation from PGi-ENK to CNA-CRF.

### Neutral trials revealed prepotent behavioral strategies

Behavioral strategies across sexes vary, with females more likely to deploy risk averse strategies (Day et al., 2016; Chowdhury et al., 2019; Kutlu et al., 2020). Behavior during neutral trials allowed us to identify rat’s underlying strategy. Females were more likely to respond during neutral trials compared to males and were more likely to respond on the avoidance lever, highlighting a critical sex difference where females prioritized safety, confirming what others have found, females use more risk averse strategies compared to males (Jolles et al., 2015; Orsini et al., 2016). Clenbuterol specifically disrupted this safety strategy on neutral trials decreasing prioritization of avoidance lever presses. This further reinforces our findings from active avoidance trials that β NE components of stress shift behavior toward passive coping or disengagement.

### Distinct noradrenergic components of stress

NE systems lie at the nexus of cue driven decision making and stress. During acute stress there is an increase in activity in the hypothalamic-pituitary-adrenal axis, LC-NE system, and correspondingly an increase in circulating levels of corticotropin-releasing factor, glucocorticoids, and NE. Depending on the behavioral measure, the impacts of acute stress can be either beneficial or detrimental. Increased central LC-NE activity during stress increases arousal which can facilitate cognitive flexibility and exploration but when more pronounced, can lead to increased habitual and disengaged behavior (Aston-Jones et al., 1997; Bouret and Sara, 2004; Schwabe and Wolf, 2009, 2011; Addicott et al., 2017; Vazey et al., 2018). This shift can contribute to the symptoms of psychological disorders such as PTSD, addiction, and depression. We show that increased NE activity on lower affinity α1 and β receptors (Arnsten et al., 2015) but not a general increase in NE tone (α2 autoreceptor inhibition (Lapiz et al., 2007)) decreased both positive and negative reinforcement behavior. Global NE increase via atipamezole did not impact positive or negative reinforcement behavior in males or females. This supports our findings that low affinity NE receptors mediate targeted effects of stress on reinforcement behavior.

### Potential peripheral impacts

Our primary interest was the impact of NE agonists on reinforcement behavior mediated by central NE signaling, however the noradrenergic component of stress has central and peripheral impacts. We used systemic administration of noradrenergic agonists that also mediated changes in blood pressure, vasoconstriction, feeding, and smooth muscle contraction through peripheral mechanisms (Civantos Calzada and Aleixandre De Artiñano, 2001; Ulrich-Lai and Herman, 2009). This includes piloerection observed in cirazoline treated animals (Supplemental Figure 1). Some observed behaviors, such as decreased movement after β agonist administration, may have been impacted by peripheral NE mechanisms. However, there was almost no effect of clenbuterol on reward seeking behavior, so it is likely that the systemic impacts were not generally inhibiting behavioral engagement.

## Conclusion

We explored the interplay between increased noradrenergic receptor activity and sex differences during reinforcement behavior. We show that males and females implement different strategies when performing a behavior to gain reward vs avoid a threat. These strategies are further evident under stress-related noradrenergic signaling. We found that females were resilient to low affinity NE signaling during reward seeking but males were not. Both males and females displayed decreased active avoidance behavior, shifting strategies to a passive coping strategy when under noradrenergic stress. These results highlight sex differences in reinforcement strategies under pharmacological stressors and provide insight into passive coping behaviors found in a number of neuropsychiatric disorders.

## CONFLICT OF INTEREST

The authors report no potential conflicts of interest.

This article was posted on a preprint server bioRxiv prior to peer review.

## ACKNOWLEDGEMENTS

This work was supported by National Institutes of Health PHS awards U01AA025481 (EMV), R00MH104716 (EMV) and F31MH131348 (EMR).

## Supplemental Figure

**Supplemental Figure 1.**
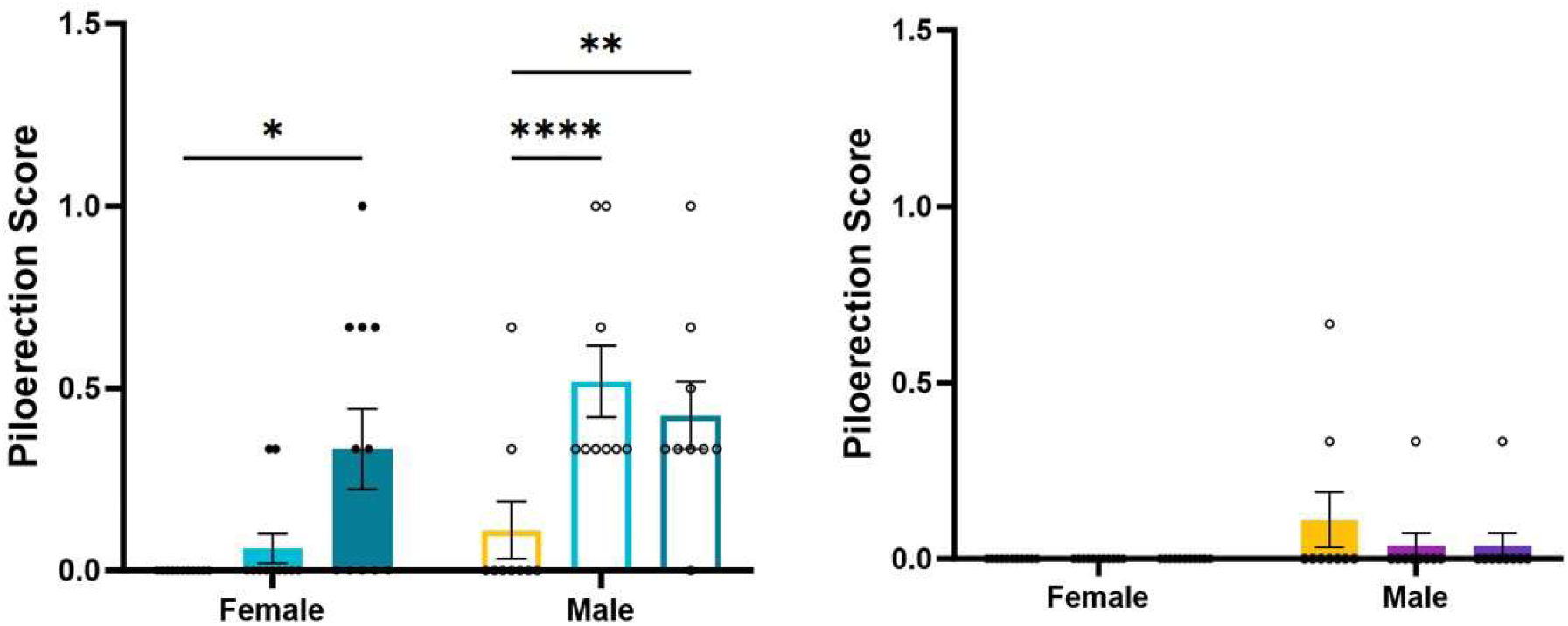
Cirazoline (left) but not Clenbuterol (right) increased physical signs of autonomous arousal, piloerection. There was an increased in experimenter identified piloerection after high doses of cirazoline in females (p=0.0231) and both dose of cirazoline increased piloerection in males (low p<0.0001, high p= 0.0039).

